# Balancing selection at a premature stop mutation in the *myostatin* gene underlies a recessive leg weakness syndrome in pigs

**DOI:** 10.1101/442012

**Authors:** Oswald Matika, Diego Robledo, Ricardo Pong-Wong, Stephen C. Bishop, Valentina Riggio, Heather Finlayson, Natalie R. Lowe, Annabelle E. Hoste, Grant A. Walling, Alan L. Archibald, John A. Woolliams, Ross D. Houston

**Affiliations:** The Roslin Institute and Royal (Dick) School of Veterinary Studies, The University of Edinburgh, Midlothian, EH25 9RG, UK; JSR Genetics, Southburn, Driffield, East Yorkshire, YO25 9ED, UK.

**Keywords:** leg weakness, MSTN, balancing selection, heterozygous advantage, pigs

## Abstract

Balancing selection provides a plausible explanation for the maintenance of deleterious alleles at moderate frequency in livestock, including lethal recessives exhibiting heterozygous advantage in carriers. In the current study, a leg weakness syndrome causing mortality of piglets in a commercial line showed monogenic recessive inheritance, and a region on chromosome 15 associated with the syndrome was identified by homozygosity mapping. Whole genome resequencing of cases and controls identified a mutation coding for a premature stop codon within exon 3 of the porcine *Myostatin (MSTN)* gene, similar to those causing a double-muscling phenotype observed in several mammalian species. The *MSTN* mutation was in Hardy-Weinberg equilibrium in the population at birth, but significantly distorted amongst animals still in the herd at 110 kg, due to an absence of homozygous mutant genotypes. In heterozygous form, the *MSTN* mutation was associated with a major increase in muscle depth and decrease in fat depth, suggesting that the deleterious allele was maintained at moderate frequency due to heterozygous advantage. Knockout of the porcine *MSTN* by gene editing has previously been linked to problems of low piglet survival and lameness. This *MSTN* mutation is an example of putative balancing selection in livestock, providing a plausible explanation for the lack of disrupting MSTN mutations in pigs despite many generations of selection for lean growth.

## Introduction

Leg weakness is heterogeneous condition causing lameness in pigs, and has negative impacts on both animal welfare and productivity [1, 2]. Significant heritability estimates have been reported for leg weakness traits [reviewed in 3], with moderate to high estimates in certain pig breeds, e.g. h^2^ = 0.45 in Landrace [4]. Several quantitative trait loci (QTL) have been identified for these traits, albeit they are generally not consistent across studies and breeds [5–8], which may be partly due to the heterogeneity of this condition. Interestingly, significant genetic correlations between leg weakness and other production traits (such as growth and muscle depth) have been detected [4]. Further, in a divergent selection experiment in Duroc lines, selection for high leg weakness was associated with a significant increase in muscle length and weight [9]. Taken together, these results suggest a degree of antagonistic genetic relationship between leg weakness and muscle growth traits in pigs, potentially explaining increases in the syndrome observed with intense selection for lean growth in recent decades.

Deleterious alleles can be maintained at relatively high frequency in commercial livestock populations due to heterozygous advantage for traits under selection [10]. Examples of such balancing selection in cattle include a frame-shift mutation in the *mannose receptor C typed 2 (MRC2)* gene responsible for crooked tail syndrome and also associated with increased muscle mass in Belgian Blue [11], and a large deletion with antagonistic effects on fertility and milk production traits in Nordic Red [12, 13] is likely to have caused an increase in incidence of porcine stress syndrome (also known as malignant hyperthermia) in the 1970s and 1980s, due to the association of the causative missense mutation with reduced backfat - a trait under selection. More recently, a recessive embryonic lethal deletion in the *bone structure and strength QTL 9 (BSS9)* gene was associated with heterozygous advantage for growth rate, explaining an unexpectedly high frequency of this allele in a commercial pig line [14].

We investigated genetic parameters and mode of inheritance for a leg weakness syndrome causing piglet mortality in a commercial line of Large White pigs. A monogenic recessive inheritance was observed, and homozygosity mapping was used to identify a genomic region on *sus scrofa* chromosome (SSC) 15 associated with the trait. A mutation causing a premature stop codon in exon 3 of the *Myostatin (MSTN)* gene (similar to mutations causing the ‘double-muscling’ phenotype in cattle [15] was the outstanding functional candidate in the region. Comparison of *MSTN* genotype frequencies at birth and 110 kg supported this hypothesis, and carriers were shown to have significantly higher muscle mass and reduced fat depth than wild type homozygotes. Therefore, we propose that the *MSTN* mutant allele is highly deleterious in this population in homozygous form, but was maintained at moderate frequency due to heterozygous advantage.

## RESULTS

### The piglet leg weakness trait shows a recessive mode of inheritance

Estimates of heritability for the leg weakness syndrome (analysed as a binary trait on the underlying liability scale) was high (0.57 ± 0.10 in the sire and dam model) with low (0.17 ± 0.02 and 0.11 ± 0.02) but significant effects observed for permanent environmental effects due to the dam and litter, respectively (Table 2, Table S1). The overall prevalence of leg weakness in the commercial cohort was 6.3 % (Table 1). When only affected litters were considered, the mean proportion of affected piglets was 23 % ± 0.7. This within-litter prevalence is consistent with the expectation under the hypothesis of a single recessive locus (i.e. 25 %). Complex Bayesian segregation analysis suggested that almost all the variation was explained by a single locus with almost no environmental variation. The estimate of the additive effect was 0.50 ± 0.001 and dominance effect was -0.50 ± 0.001, which is in precise agreement with a recessive locus model.

**Table 1.** Number of animals in pedigree and records used for variance components analyses

| Description | Data |
| --- | --- |
| Number of records | 19,006 |
| Number of pedigree records | 27,501 |
| Number of generations | 9 |
| Number of litter | 1,903 |
| Number of Sires | 346 |
| Number of Sires of Sire | 175 |
| Number of Dams of Sire | 239 |
| Number of Dams | 1,929 |
| Number of Sires of Dam | 240 |
| Number of Dams of Dam | 882 |
| Prevalence % | 6.3 |

**Table 2.** Genetic parameter estimates and standard errors for the trait of leg weakness on the liability scale using the logit transformation showing the outcomes of fitting sire or dam models with or without maternal environment (σ^2^_w_). All models have litter variance (σ^2^_v_) fitted

| Model<sup>¥</sup> | Sire | Sire and Dam Models |  |
| --- | --- | --- | --- |
| | $\sigma^2_w > 0$ | $\sigma^2_w > 0$ | $\sigma^2_w = 0$ |
| $\sigma^2_s$ | 1.069 (0.283) | 1.100 (0.290) | 1.087 (0.286) |
| $\sigma^2_d$ | | 0.664 (0.231) | 1.110 (0.186) |
| $\sigma^2_v$ | 0.681 (0.123) | 0.681 (0.123) | 0.769 (0.123) |
| $\sigma^2_w$ | 0.999 (0.148) | 0.437 (0.203) | |
| $\sigma^2_p$ | 6.049 (0.307) | 6.182 (0.322) | 6.267 (0.325) |
| $h^2_u$ | 0.71 (0.16) | 0.57 (0.10) | 0.70 (0.08) |
| $\sigma^2_v/\sigma^2_p$ | 0.11 (0.02) | 0.11 (0.02) | 0.12 (0.02) |
| $\sigma^2_w/\sigma^2_p$ | 0.17 (0.2) | 0.07 (0.03) | |

<sup>¥</sup> - $\sigma^2_s$ , $\sigma^2_d$ , $\sigma^2_v$ and $\sigma^2_w$ are variances due to the sire and dam genetic effects, litter and maternal environment effects. $\sigma^2_p$ is phenotypic variance. $h^2_u$ is estimated heritability on observed scale, and $\sigma^2_v/\sigma^2_p$ and $\sigma^2_w/\sigma^2_p$ are the proportion of phenotypic variance explained by the litter and maternal environment effects respectively

### Causative variant maps to chromosome 15

Homozygosity mapping was used to map the underlying recessive variant, and the longest homozygous segment was a region of ~ 8.3 Mbp on SSC 15. In this region, all 55 informative single nucleotide polymorphisms (SNPs) on the Illumina PorcineSNP60 SNP chip [16] were homozygous in affected animals, but contained SNPs that were heterozygous or homozygous for the alternative allele in the unaffected animals. The segment started with SNP ALGA0110636 (rs81338938) at position 86,745,668 in the current Sscrofa11.1 reference genome assembly (Genbank assembly accession GCA_000003025.6) and finished with SNP H3GA0044732 (rs80936849) at position 95,062,143 (Fig 1a and b). The *MSTN* gene was located within this homozygous segment, from position 94,620,269 - 94,628,630. The 8.3 Mbp segment was assumed to represent a selective sweep likely to contain the underlying causative mutation, and became the focus of further analyses to discover and characterise this mutation.

**Fig 1.**
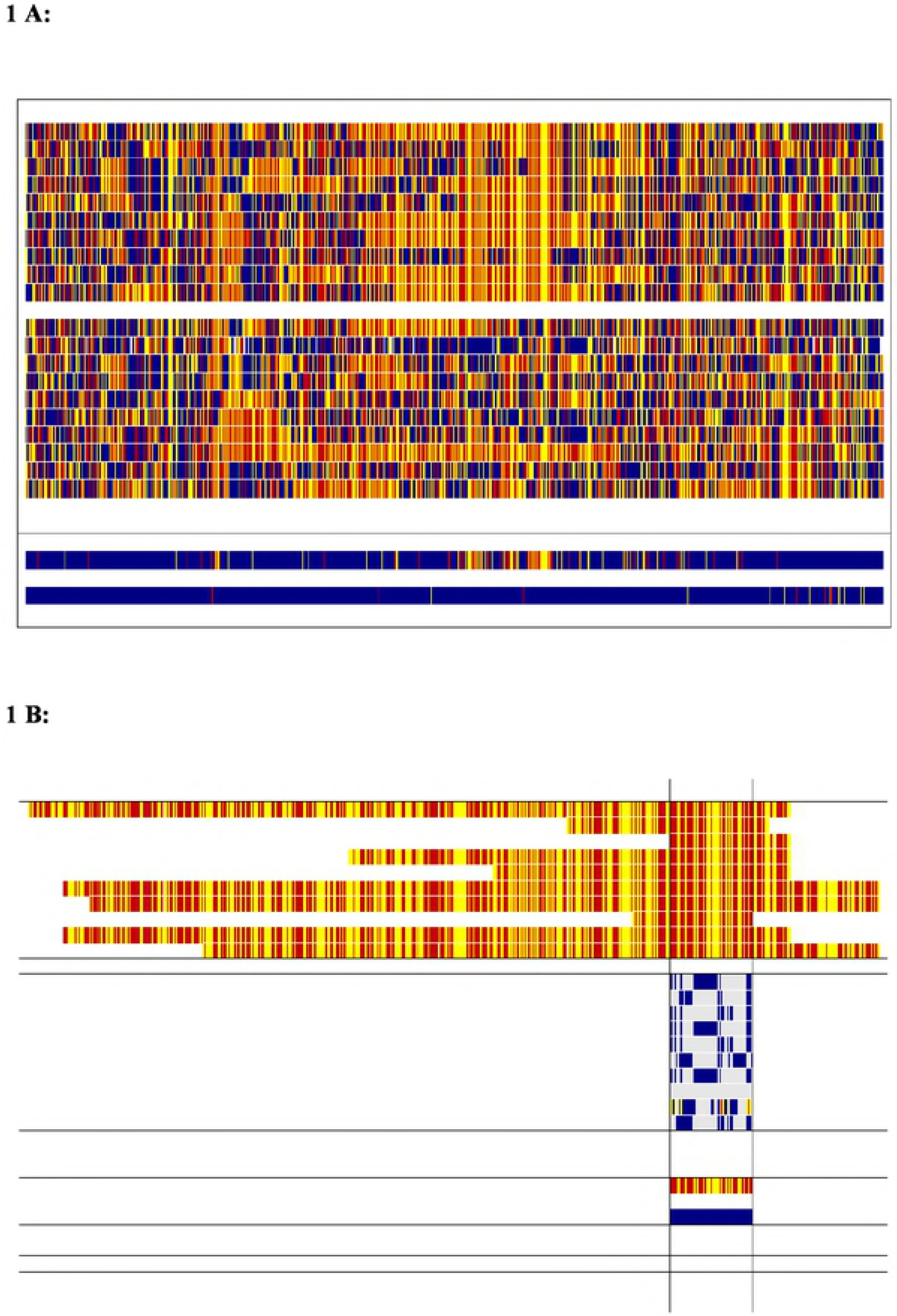
**A:**Homozygosity mapping of the leg weakness syndrome assuming a single underlying recessive mutation on Chromosome 15 (SSC15). Vertical blocks in red and yellow represent homozygous genotypes, and blue the heterozygote genotype. The ten cases (above) and ten controls (below) are shown (one per line). A summary of homozygosity mapping is provided cases vs control at the bottom of Fig1A in which if all the animals within a group (i.e. case or control) are homozygous for the same allele, then the relevant colour (red or yellow) is shown and if any animal within a group (i.e. case or control) is heterozygous then the SNP is coloured blue **B:** This is an extract from Fig1a, showing the longest shared haplotype segment (55 SNPs) in the cases on SSC15 ranging from ALGA0110636 (rs81338938) to H3GA0044732 (rs80936849) and corresponds to position 86,745,668– 95,062,143 in the new pig reference genome assembly Sscrofa11.1 GCA_000003025.6). The first ten lines are the cases; the second ten lines are the controls with genotypes shown only for the region of homozygosity shared across the controls. Genotypes shown for controls are shown only if they are different to cases (blank genotype in controls within the targeted segment means that they share the same genotype as cases). Finally, the bottom two lines are summary lines shown on the same basis as the summary lines in Fig1A.

### Sequence analysis reveals *MSTN* mutation as causative candidate

To identify candidates for the causative variant, whole genome sequence data from ten cases, six presumed heterozygous carrier dams, and 22 controls were analysed. A total of 40 SNPs identified within the homozygous segment fitted the pattern of a potential causative variant assuming a recessive mode of inheritance. Functional annotation of these SNPs revealed that 19 were intergenic, 19 were intronic, 1 was in a pseudogene, and 1 caused a premature stop codon. There were also 10 InDels identified, 3 of which were intergenic and 7 of which were intronic (Figure S1). The outstanding functional candidate was a mutation in the third exon of the *MSTN* locus that results in the replacement of a codon for glutamic acid with a stop codon in exon 3 at position 274 (c.820G>T; p.E274*) (Figure 2A). The mutation is located in a region that is highly conserved across multiple species, and is predicted to result in truncation of the protein (Figure 2B). Functional annotation of all other variants detected in the selective sweep region did not reveal any other obvious causative candidates. This SNP and stop gain mutation was not present on the Ensembl variation database, accessed 25^th^ July 2018.

**Fig 2.**
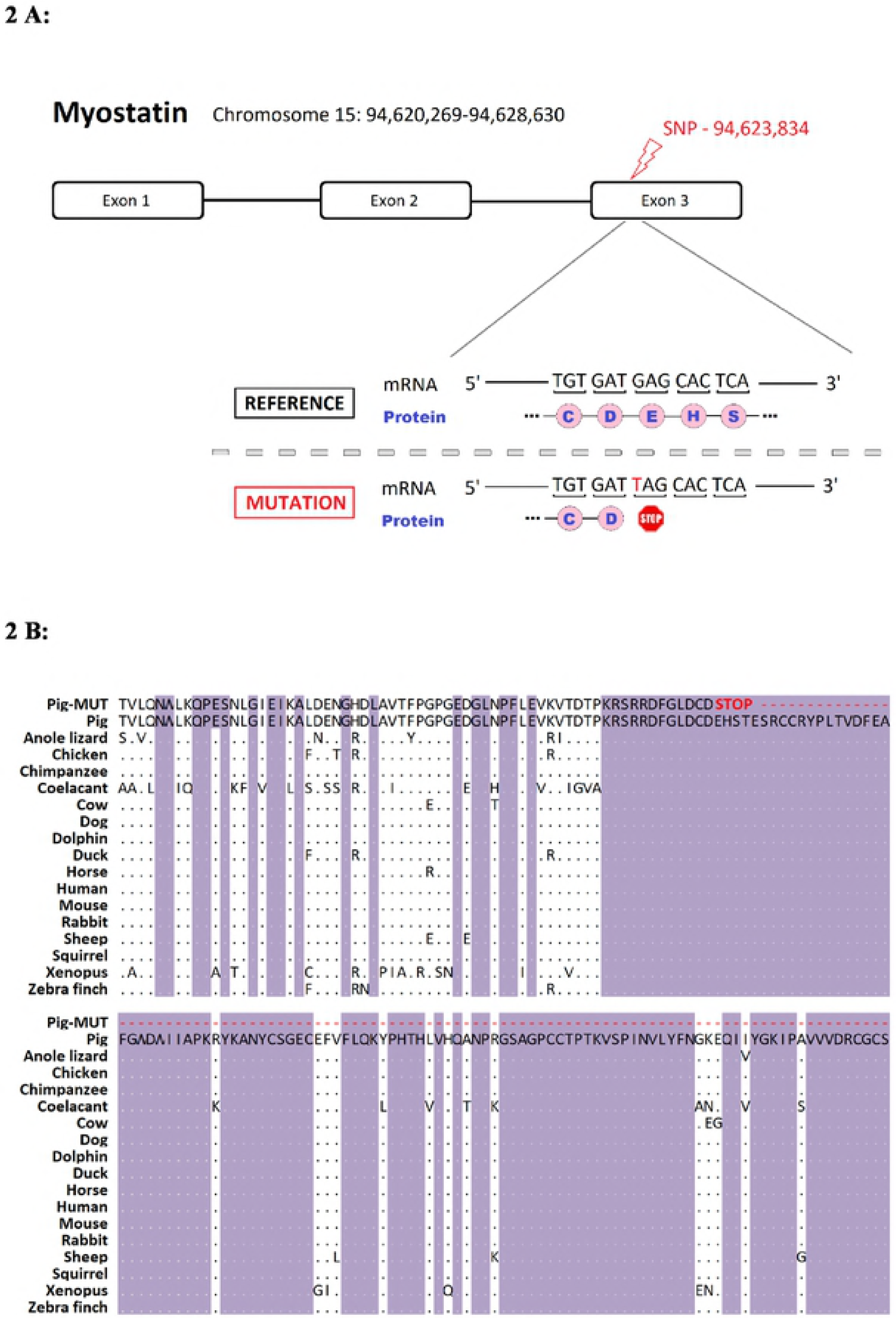
(A) Position of the premature stop causing mutation within the porcine myostatin locus; (B)Conservation of the amino acid sequence surrounding the mutation, with the consequences of the premature stop mutation highlighted in red.

### Changes in mutant allele frequency from piglets to adults

The *MSTN* c.820G>T mutation showed no statistically significant deviation from Hardy Weinberg equilibrium (HWE) in the 486 piglets sampled at birth (q = 0.22, α = 0.019, χ^2^=0.18, P > 0.05). Random mating of the dioecious population would be expected to result in a value of α that is slightly negative [17], and the value observed does not differ significantly from this value. However, the mutation deviated significantly from HWE at 40 kg (q = 0.17 α = - 0.180, χ^2^=12.2, P > 0.001) and at 110 kg (q = 0.17 α = - 0.210, χ^2^=11.45, P > 0.001). This was due to the loss of homozygous mutant piglets, with all but one dying (or being euthanized) shortly after birth, and the remaining piglet being euthanized due to poor health before it reached 110 kg. The large change in α in a negative direction is quantitative evidence of the selective disappearance of homozygote genotypes, as opposed to disappearance as a result of selection against the allele itself. There were no significant changes in the relative genotype frequencies (GG, GT) over the total period or any sub-period from birth to the end of the test, confirming all changes in *q* and *a* are due to the selective loss of homozygotes, and that any other mortality or culling was at random with respective to *MSTN* genotype.

### Association of the *MSTN* mutation with performance traits

The association of the porcine *MSTN* c.820G>T mutation with performance traits was assessed on 384 pigs which had completed a commercial performance test. Given the loss of the homozygous mutant animals the effect of the *MSTN* c.820G>T mutation was only estimated by the difference between the heterozygotes and the wild type pigs. The genotype means and differences are shown in Table 3, with the most notable of these being a major increase in muscle depth and a reduction in fat depth in the carriers (p < 0.001), with no evidence of a difference in live weight at 110 kg. Approximately 31 % of the genetic variation in muscle depth and 18 % of the genetic variation in fat depth was explained by this single variant (Table 3). The heterozygous animals had on average 5 mm increased muscle depth, and 1.7 mm decreased backfat depth when compared with wild type homozygous animals.

**Table 3.**
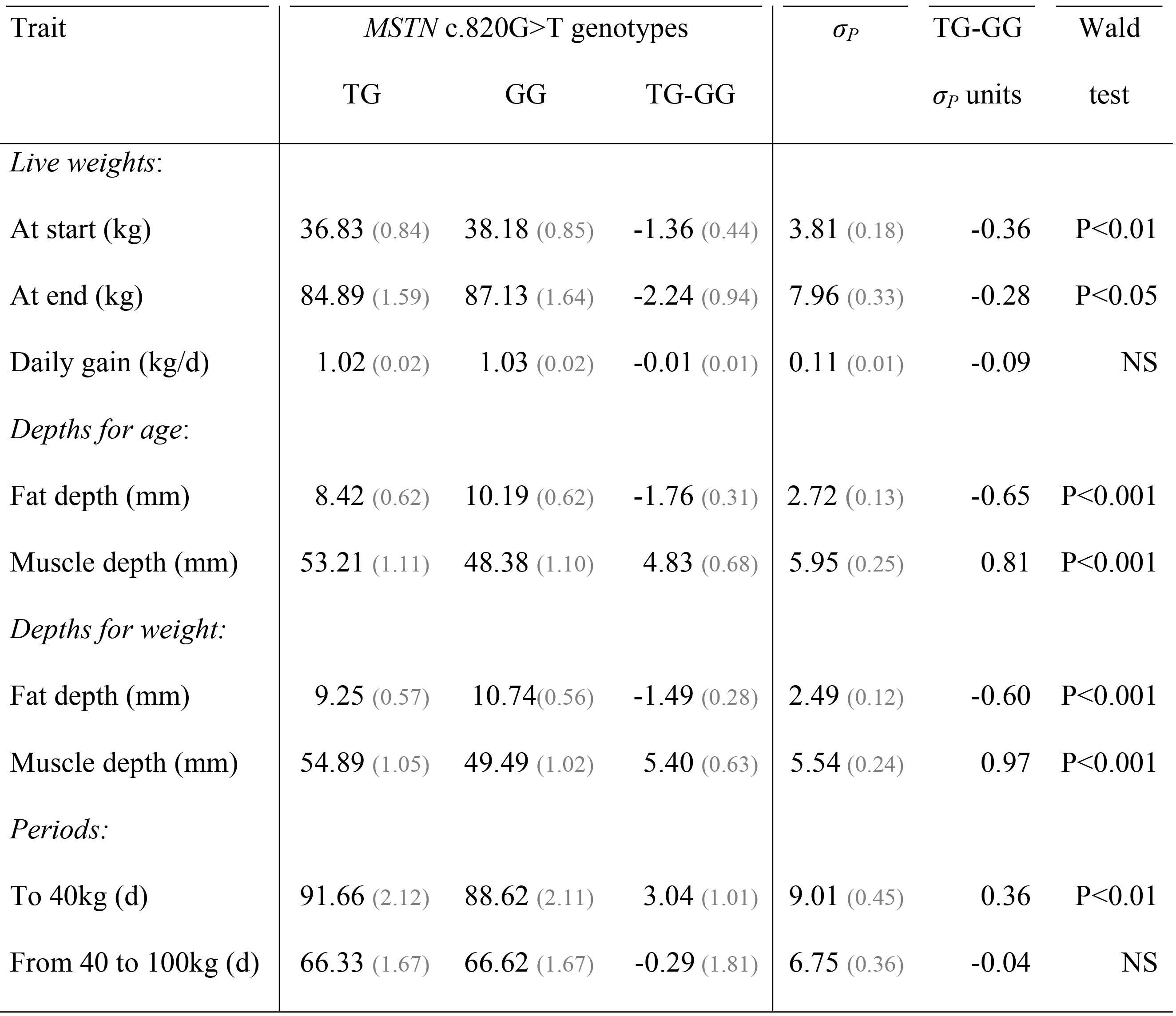
Estimates and statistical significance of the effect of the *MSTN* c.820G>T locus on the growth and carcass traits of pigs obtained from a commercial performance test. Estimates are shown in absolute units and standardised by phenotypic standard deviations (*σ*_*p*_). Standard errors are in parentheses. The traits are categorised into: live weights and live weight gain; muscle and fat depths measured by ultrasound at the end of the test either conditional on age or on live weight; and periods to achieve growth targets.

| Trait | *MSTN* c.820G>T genotypes | | | $\sigma_P$ | TG-GG | Wald |
| --- | --- | --- | --- | --- | --- | --- |
| | TG | GG | TG-GG | | $\sigma_P$ units | test |
| *Live weights:* |  |  |  |  |  |  |
| At start (kg) | 36.83 (0.84) | 38.18 (0.85) | -1.36 (0.44) | 3.81 (0.18) | -0.36 | P<0.01 |
| At end (kg) | 84.89 (1.59) | 87.13 (1.64) | -2.24 (0.94) | 7.96 (0.33) | -0.28 | P<0.05 |
| Daily gain (kg/d) | 1.02 (0.02) | 1.03 (0.02) | -0.01 (0.01) | 0.11 (0.01) | -0.09 | NS |
| *Depths for age:* |  |  |  |  |  |  |
| Fat depth (mm) | 8.42 (0.62) | 10.19 (0.62) | -1.76 (0.31) | 2.72 (0.13) | -0.65 | P<0.001 |
| Muscle depth (mm) | 53.21 (1.11) | 48.38 (1.10) | 4.83 (0.68) | 5.95 (0.25) | 0.81 | P<0.001 |
| *Depths for weight:* |  |  |  |  |  |  |
| Fat depth (mm) | 9.25 (0.57) | 10.74(0.56) | -1.49 (0.28) | 2.49 (0.12) | -0.60 | P<0.001 |
| Muscle depth (mm) | 54.89 (1.05) | 49.49 (1.02) | 5.40 (0.63) | 5.54 (0.24) | 0.97 | P<0.001 |
| *Periods:* |  |  |  |  |  |  |
| To 40kg (d) | 91.66 (2.12) | 88.62 (2.11) | 3.04 (1.01) | 9.01 (0.45) | 0.36 | P<0.01 |
| From 40 to 100kg (d) | 66.33 (1.67) | 66.62 (1.67) | -0.29 (1.81) | 6.75 (0.36) | -0.04 | NS |

## DISCUSSION

A piglet leg weakness syndrome identified in a commercial line of Large White pigs showed moderate to high heritability, consistent with the upper range of estimates reported in the literature [18, 19, 2]. The within-litter incidence of the syndrome and the complex Bayesian segregation analysis both pointed to a monogenic recessive condition. Homozygosity mapping revealed a 8.3 Mbp segment on SSC15 as a putative selective sweep likely to contain the causative variant. The outstanding functional candidate variant identified by whole genome sequencing of cases and controls caused a premature stop codon within exon 3 of the *MSTN* gene (Figure 2), and this variant was in HWE at birth but significantly distorted by 110 kg due to an absence of homozygous mutant animals. Knockout of *MSTN* by gene editing in previous studies has been associated with poor health and mortality of knockout piglets [20–23], including observations of a piglet leg weakness syndrome with an inability to stand or walk [21], strikingly similar to the syndrome described in the current study.

*MSTN* is a member of the *transforming growth factor beta* (*TGF-β*) superfamily, which is highly conserved across species, and is typically expressed in developing and mature skeletal muscle as a key regulator of muscle growth [24]. The *MSTN* gene has been a gene of interest to animal breeders for over twenty years since the discovery of loss-of-function mutations in the cattle *MSTN* gene, which cause muscle hypertrophy leading to double muscling phenotypes [15, 24, 25]. Interestingly, these loss-of-function mutations causing double muscling are frequently due to premature stop mutations in the highly conserved exon 3 of *MSTN [25]*, (Figure 3). Further, in the case of Marchigiana beef cattle, the *MSTN* variant causing double muscling results is due to replacement of Glutamic Acid with a stop codon (the same change observed in the current study) in the equivalent position in the protein as observed in the current study [26]. This observation provides additional indirect evidence that the porcine *MSTN* mutation described herein is likely to cause the increase in muscle depth and decrease in fat depth associated with carrier animals.

In livestock breeding schemes, *MSTN*-inactivating mutations have been retained due to selection for increased lean growth associated with meat production, particularly for double-muscled cattle. However, despite many generations of selection for lean growth in pigs no such mutations have been reported previously. Associations between polymorphisms within the porcine *MSTN* gene and production traits have been shown in small-scale studies [27–29]. Further, a genome-wide association study in a sire line Large White population (related to the animals in the current study) detected SNPs on SSC 15 significantly associated with rib fat between 81.1 and 87.8 Mbp [30]. While these SNPs are between 7 and 13 Mbp closer to the centromere than *MSTN*, two of these SNPs overlap with the region on homozygosity indicative of a selective sweep in the current study. Therefore, it is plausible that this association may be due to linkage disequilibrium (LD) with the *MSTN* variant, or that other variants impacting performance traits exist on SSC15. The selection index applied in this line explicitly benefited animals with positive muscle depth and negative fat depth estimated breeding values, with analysis of the index showing that heterozygotes had a slight but significant advantage (p<0.05). Therefore, it is likely that the MSTN c.820G>T mutation was maintained at moderate frequency in this line despite its deleterious impact on piglet mortality due to heterozygous advantage for muscle and fat traits. Similar examples of balancing selection have observed to explain the maintenance of deleterious alleles at moderate frequency in commercial cattle [10, 31] and pig [14] populations. The results described herein have major implications for the targeted ablation of *MSTN* via gene editing to increase lean growth in pigs, and provide a plausible explanation of why *MSTN* loss-of-function mutations have not previously been reported in pigs despite decades of selection for lean growth.

## Materials and Methods

### Ethics statement

All samples were collected on a commercial nucleus farm as part of standard husbandry and management procedures in the nucleus herd, which complied with conventional UK red tractor farm assurance standards (https://assurance.redtractor.org.uk/) where sick or injured livestock that do not respond to treatment are promptly and humanely euthanized by a trained and competent stockperson.

### Animals

A piglet leg weakness syndrome was characterised in a Large White sire line, reared in a nucleus herd, under standard conditions but with additional data recording. Leg weakness trait observations were collected on 19,006 piglets from 2007 to 2010, during which time a high incidence of the syndrome was detected. In addition, DNA was sampled on a further 119 piglets in 2011 and 486 piglets from the same population in 2012, of these 384 also had weight and carcass phenotypes available. The cohort born in 2007 to 2010 will be referred to as the commercial cohort and those in 2011 and 2012 as the survey cohort. Pedigree was available for all animals and spanned 9 generations. Details of the data and pedigree structure are presented in Table 1.

The leg weakness in the phenotyped animals was visually classified as normal or affected (0 / 1 respectively). The leg defect is characterised by the piglet not being able to straighten its legs to stand, this being most apparent for the front legs, and being slow to suckle. Videos exemplifying the syndrome are given in Additional File 1. Detailed post mortems were conducted on two affected individuals but the results were ineffective in providing additional diagnostic aids (see Additional File 2). The Online Mendelian Inheritance in Animals database (OMIA: http://omia.org/0MIA000585/9823/) was searched for previous reports of leg weakness in pigs but the syndrome observed here did not appear. These problems frequently resulted in death from either starvation or being crushed by the sow. Where the outcome resulted in poor welfare the piglets were euthanised in accord with the Ethics Statement. Additional farrowing data were collected on the females in the commercial cohort including numbers born alive, dead or mummified, parity and year of birth. Body weights, ultrasound muscle and fat depths were obtained for individuals in the survey cohort that were retained in the herd until commercial slaughter age (when average weight is 110kg). The ultrasonic measurements taken were the average depth of the *m. longissimus dorsi* and overlaying subcutaneous fat layer across the last four ribs.

### Statistical analyses of genetic parameters

Initial studies to establish the genetic basis of the leg weakness syndrome were undertaken by fitting linear mixed models to the commercial cohort. The binary record of the syndrome was modelled on the observed 0 / 1 scale and the full model fitted was:

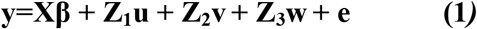

where **y** is the vector of leg weakness phenotypes; **β**, a vector of fixed effects for month of observation (43 df), sex (1 df), parity (7 df), numbers in litter born dead (1 df) or alive (1 df), with design matrix **X**; **u**, additive polygenic effects, assumed to be distributed MVN(0, **A**σ^2^_a_) with design matrix **Z_1_**; **v**, random litter effects assumed to be distributed MVN(0, **I**σ^2^_v_), with design matrix **Z**_2_; **w**, maternal environment effects across litters assumed to be distributed MVN(0, **I**σ^2^_w_) with design matrix **Z**_3_; and **e** residuals assumed to be distributed MVN(0, **I**σ^2^_e_). Variations in this model were fitted replacing the individual polygenic effects with sire and/or dam effects, assumed to be distributed MVN(0, **A**σ^2^_s_) and MVN(0, **A**σ^2^_d_) respectively. All models were fitted using the ASREML software [32]. Likelihood ratio tests were used to assess the random effects. In addition, an analogous threshold mode with an underlying continuous liability was fitted with a logit link function and sire and dam effects associated with the pedigree, but not with individual polygenic effects, following recommendation of Gilmour et al. [36]. For the full model, the phenotypic variance was calculated as σ^2^_p_ =σ^2^_u_ + σ^2^_v_+ σ^2^_w_ + σ^2^_e_. Where sire and dam models were used, σ^2^_u_ was replaced by σ^2^_s_ + σ^2^_d_. Heritability (h^2^) was calculated as σ^2^_u_/σ^2^_p_ or 2(σ^2^_s_ + σ^2^_d_)/σ^2^_p_ depending on the model. The proportion of variance explained by the litter and maternal environmental effects were estimated as σ^2^_v_/σ^2^_p_ and σ^2^_w_/σ^2^_p_, respectively. Heritabilities on the observed scale (0/1) was transformed to an underlying liability scale following Dempster and Lerner [33] using the observed prevalence of the syndrome in the commercial population which was 6.3%

Inspection of the data suggested that the syndrome may be due to a single gene with the predisposing deleterious allele showing a recessive mode of inheritance, and this hypothesis was tested using chi square tests and segregation analyses. An initial test of a monogenic recessive mode of inheritance was carried out by pooling all affected litters, estimating the probability of being affected conditional on being born in an affected litter, and using chi-squared to test the null hypothesis that the probability of being affected was 0.25. A weakness of this approach is that some litters by chance will have no affected offspring, so a more complex segregation model was fitted. This model included all known phenotypes and pedigree data, and assumed a monogenic inheritance with environmental variation fitted by Gibbs sampling [34].

### Homozygosity mapping of the recessive mutation

Ten affected animals from different litters and 10 unaffected full-sib controls from the commercial cohort were genotyped using the Illumina PorcineSNP60 SNP chip [16]. Only those SNPs that mapped to known positions on autosomal chromosomes and were not fixed nor completely heterozygous were retained. This left 38,570 segregating autosomal SNPs for the use in homozygosity mapping. Homozygous regions were assessed by alignment with the Sscrofa11.1 reference genome assembly sequence (Genbank assembly accession GCA_000003025.6).

### Whole genome resequencing

The genomes of the ten cases used for homozygosity mapping and six separate dams with affected offspring, assumed heterozygotes, were whole genome shotgun sequenced on an Illumina HiSeq 2500 platform. The dams were individually sequenced with a 10x genome coverage. The piglets were barcoded and individually sequenced at 3x coverage to achieve 30x coverage for the pool. The full sequencing output resulted in ~1.3 billion paired-end reads with an average of 48 million paired-end reads/sample for the piglets and 157 million paired-end reads/sample for the dams. Quality filtering and removal of residual adaptor sequences was conducted on read pairs using Trimmomatic v.0.32 [35]. Only reads where both pairs had a length greater than 32 bp post-filtering were retained, leaving a total of ~1.2 bn paired-end reads.

Whole genome resequencing was followed by alignment to the Sscrofa11.1 assembly; using the Burrows-Wheeler Aligner with default parameters [36]. The average alignment rate of properly paired reads was of 92 %. PCR duplicates were marked using Picard Tools (http://broadinstitute.github.io/picard). Variant calling was performed using the Genome Analysis Toolkit (GATK) HaplotypeCaller after read recalibration [37]. The parameter setting for the hard filters that were applied to the raw genotypes were: QualByDepth < 2.0, FisherStrand > 60.0, RMSMappingQuality < 40.0, MappingQualitySumTest < -12.5, ReadPosRankSumTest < -8.0.

Candidate loci were identified from the sequences of the 16 animals from this study plus 22 additional *Sus scrofa* control sequences obtained from a public database [38], comprising 7 domesticated breeds (Duroc, Hampshire, Jiangquhai, Landrace, Large White, Meishan and Pietrain) and wild boar. The following criteria were used to identify candidate SNPs:

i. Homozygous for the same allele in all the affected piglets
ii. Heterozygous in parents of affected piglets (i.e. putative carriers)
iii. Heterozygous or homozygous for the alternative allele (i.e. the one not observed in the affected offspring) in the control (unaffected) animals

It is worth noting that the limited sequencing depth means that both alleles will not be detected for all bases in all individuals. This limitation is particularly relevant to the reliable detection of heterozygous SNPs.

### Genotyping

A ‘kompetitive allele specific PCR’ (KASP) assay was designed by LGC Genomics (Teddington, UK) to enable genotyping of the mutation in the *MSTN* stop codon in large numbers of animals. The survey cohort of 486 piglets sampled at birth were genotyped by LGC. Of these, 265 remained as candidates for the final selection at 110 kg with a complete record of their performance test, together with another 119 pigs phenotyped at slaughter age (total n = 374). In both the 486 piglets and the subsequent subsets surviving to 40 and 110 kg, the frequency of the *MSTN* mutation (q) was calculated by counting, and the departure from Hardy Weinberg equilibrium the genotypes was estimated as α =1-H_obs_/H_exp_ [17] where H_obs_ is the observed heterozygosity and H_exp_ is the expected heterozygosity calculated as 2q(1-q). The significance of departure from true random mating genotype frequencies (α = 0) was tested using a chi-squared test.

### Association analysis

Associations between the *MSTN* c.820G>T locus and variation in performance traits in commercial testing conditions were examined in the survey cohort (n = 384). The performance test was started at a target weights of 40 kg, at an average age of 85 days, and continued for 54 (s.d. 12) days. The performance tests were performed over two distinct periods. The traits available were live weights and ages at the start and the end of the test, ultrasonic muscle and fat depths measured at the end of the test, days from birth to 40 kg and days from 40 to 110 kg. Univariate mixed models were fitted to these data in ASReml-R4 using the following model:

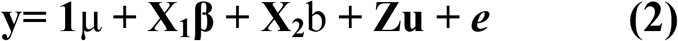

where **y** is the vector of phenotypes; *μ*, a fitted mean, and **1**, a vector of 1’s; **β**, a vector of fixed nuisance effects with design matrix **X_1_**; *b*, a scalar fixed effect for the effect of SNP genotype with design matrix **X_2_**, this has only 1 df due to the absence of homozygotes completing the test; **u**, additive polygenic effects assumed to be distributed MVN(0, **A**σ^2^_a_), with design matrix **Z**; and **e**, residuals assumed to be distributed MVN(0, **I**σ^2^_e_). For all traits, the sex of the piglet (1 df), parity of dam (4 df) and period of testing (1 df) were fitted as nuisance factors, together with cubic smoothing splines for the start date of the test fitted separately within each period [39]. The age at the time of measurement was fitted as covariate (1 df) for all traits other than days to 40 kg and days from 40 to 110 kg. The significance of fixed effects was assessed using Wald tests [ASReml Manual].

## ACKNOWLEDGEMENTS

Institute Strategic Funding Grants to The Roslin Institute (BBS/E/D/20211553 BBS/E/D/30002275). Edinburgh Genomics is partly supported through core grants from NERC (R8/H10/56), MRC (MR/K001744/1) and BBSRC (BB/J004243/1). Diego Robledo is supported by a Newton International Fellowship of the Royal Society (NF160037)

